# *Klebsiella pneumoniae* isolated from the intestines of *Tenebrio molitor* larvae (Coleoptera: Tenebrionidae) that consume expanded polystyrene

**DOI:** 10.1101/2024.09.20.613978

**Authors:** Luis Caruajulca-Marin, Katherin Huaman-Ventura, Marilin Sanchez Purihuaman, Junior Caro-Castro, Ada Barturen-Quispe, Segundo Vasquez-Llanos, Carmen Carreno-Farfan

## Abstract

Plastics such as polystyrene are resistant to biodegradation, pollute the environment, and negatively impact the health of living organisms. However, several organisms, such as the larvae of *Tenebrio molitor* (Coleoptera: Tenebrionidae) and their associated gut microbiome, contribute to its degradation. The aim of this research was to determine the efficiency of expanded polystyrene (EPS) degradation by gut bacteria isolated from *T. molitor* larvae. To achieve this, a set of EPS-degrading bacteria was selected based on the time required to utilize the polymer as a carbon and energy source. Additionally, EPS degradation efficiency was compared, and the most efficient degrading bacterium was identified at the molecular level. Results showed that 95.13% of the bacteria isolated on nutrient agar and 86.57% of those isolated on McConkey agar were able to grow on EPS, with five bacteria being selected that utilized the polymer after 36 hours of incubation. The efficiency of EPS degradation, expressed as the percentage of weight loss by the degrading bacteria, ranged from 5.29% to 12.68%, with a reduction rate of 0.0005 to 0.0013 g per day and a half-life of 533.15 to 1386.20 days. Finally, 16S rRNA gene analysis identified the bacterium as *Klebsiella pneumoniae*. Cultivable gut bacteria from *T. molitor* larvae have demonstrated potential as candidates for EPS degradation, and biotechnological techniques can further enhance the efficiency of the degradation process.

## 1. INTRODUCTION

Petroleum-derived plastics, or petroplastics, are components of most everyday materials due to their lightweight, durability, and low cost. As a result, global production has increased exponentially, reaching over 390.7 million metric tons (MT) per year, compared to 1.5 million MT produced in 1950 (Shukla *et al*., 2024; United States Agency for International Development [USAID], 2020). Most plastics are single-use and do not degrade easily, leading to an estimated 4.9 billion MT discarded between 1950 and 2005 that persist somewhere in the world (Song *et al*., 2020). Petroplastics in the form of macroplastics (>25 mm), mesoplastics (5-25 mm), large microplastics (1-5 mm), small microplastics (20 μm-1 mm), and nanoplastics (1-1000 nm) contaminate soil, water, and air, impacting the health of all living beings (Shukla *et al*., 2024; Hwang *et al*., 2020).

Polystyrene (PS) is a polymer with an amorphous structure, high rigidity, and lightweight properties, making it low-cost and fully recyclable. However, the recycling market is not significant. Additionally, it is marketed in various forms, including expanded polystyrene (EPS) (Song *et al*., 2020). Due to poor segregation of EPS, it is found contaminating various levels of the ecosystem. Oceans contain high amounts of EPS, which, when exposed to sunlight, fragments and generates microplastics and nanoplastics (Du *et al*., 2024). PS has also been detected in soil at depths of 0-30 cm in countries such as Germany, China, and Spain (Dioses *et al*., 2020). Furthermore, PS has been reported as an air pollutant in studies conducted in Paris, London, and Dongguan (Amato *et al*., 2020). In Peru, pollution from plastics such as PS (De la Torre *et al*., 2024) is increasing daily, with more than 500 microplastics per square meter recorded on the beaches of Lima and Callao (USAID, 2020).

In living organisms, the presence of PS in the soil has been shown to alter photosynthesis (Zhuang *et al*., 2024), reduce chlorophyll (Iswahyudi *et al*., 2024), carotenoids, micronutrients, as well as plant growth and biomass (Lian *et al*., 2021). In animals, PS can reduce sperm quality and serum reproductive hormones (Ying *et al*., 2024), alter ovarian function, decreasing oocyte quality and fertility (Xue *et al*., 2024), and cause dysregulation of genes associated with neurotransmitters in embryos (Shubham *et al*., 2024). Additionally, it can increase enzymes related to oxidative stress (Zhen *et al*., 2024), cause toxicity in hemolymph and digestive glands (Ventura *et al*., 2024), and increase alcohol/aldehyde dehydrogenase (AADH) and lactate dehydrogenase (LDH) enzymes in tissues, reducing survival (Gagné *et al*., 2023). On the other hand, it can reduce the rate of photosynthesis and nitrogen fixation in cyanobacteria (Lixia *et al*., 2024).

Humans ingest styrene monomers from various sources such as food, water, medical devices, and everyday products. It has been estimated that human consumption of PS can exceed 133 mg per year when particles have a diameter of 100 μm (Hwang *et al*., 2020). PS nanoparticles (25-70 nm in diameter) can be absorbed by living organisms, significantly affecting the viability of human alveolar epithelial cells, activating genes related to the inflammatory response, modifying the expression of proteins associated with the cell cycle, and inducing pro-apoptosis (Xu *et al*., 2019).

In nature, various larval forms of insects, such as *Tenebrio molitor* Linnaeus, 1758 (Yang *et al*., 2023), *T. obscurus* (Peng *et al*., 2019), *Uloma* sp. (Kundungal *et al*., 2021), *Tribolium castaneum* (Rodríguez *et al*., 2021), *Zophobas atratus* (Lu *et al*., 2024), *Z. morio* (Sun *et al*., 2022), *Galleria mellonella* (Young *et al*., 2024), and *Alphitobius diaperinus* (Cucini *et al*., 2022), are known to fragment polystyrene (PS) to consume and ingest it into their intestines. In the intestines, PS is ultimately mineralized by specialized intestinal bacteria, including *Acinetobacter, Alcaligenes, Bacillus, Citrobacter, Enterobacter, Klebsiella, Massilia, Pseudomonas*, and *Serratia* (Sufang *et al*., 2024; Machona *et al*., 2022; Vera *et al*., 2021; Tan *et al*., 2021a; Urbanek *et al*., 2020; Jiang *et al*., 2021; Brandon *et al*., 2021). These bacteria degrade the polymer into carbon dioxide and water, presenting a potential alternative for reducing this pollutant (Wang *et al*., 2020; Peng *et al*., 2019). Therefore, the objective of this study was to determine the efficiency of expanded polystyrene degradation by cultivable bacteria associated with the intestines of *Tenebrio molitor* larvae (Coleoptera: Tenebrionidae).

## 2. MATERIALS AND METHODS

### 2.1. Preparation of emulsified EPS for feeding *T. molitor* larvae

The *T. molitor* larvae (20-24 mm) were purchased from the online store Kuru Wasi in Lima, Peru, while the EPS was obtained from ETSAPOL, SKU: 11760. To prepare the emulsified EPS, 3 g of EPS were placed in a beaker and dissolved in 97 mL of dichloromethane (Spectrum), then allowed to rest at 4°C for 48 hours. The emulsified EPS sheets adhered to Petri dishes were carefully removed, cut into 1 cm^2^ fragments, and placed in tubes with pre-sterilized carbon-free minimal basal broth (Tan *et al*., 2021b). The wheat bran was sterilized at 120°C for 10 minutes and then dried at 80°C for 10 minutes, while the EPS was sterilized at 80°C for 10 minutes. Both were cooled to room temperature before being fed to the larvae (Bae *et al*., 2021).

### 2.2. Quantification of cultivable bacteria associated with the intestinal tract of *T. molitor* larvae

The number of colony-forming units (CFU mL^-1^) of bacteria from the intestinal content was determined in 60 *T. molitor* larvae sourced from three different conditions: directly from the commercialization center (LCC), from a laboratory rearing with a mixed diet of wheat bran and EPS (LRMD), and from a rearing with a diet consisting solely of EPS (LREPS) over a period of 21 days (Tan *et al*., 2021a). The larvae (20 from LCC, 20 from LRMD, and 20 from LREPS) were first chilled at -4°C for 15 minutes to inactivate them (Vera *et al*., 2021). They were then disinfected by immersion in 75% ethanol for 1 minute, followed by rinsing twice with 0.85% NaCl solution (p/v) (Tan *et al*., 2021a) and dried with sterile filter paper. Each larva was placed on a Petri dish, and pressure was applied to the anal area to extract the intestines, which was then placed in a tube with 1 mL of 0.85% NaCl solution (p/v). Decimal dilutions up to 10^−4^ were prepared from this tissue, and these dilutions were plated in triplicate on Plate Count agar to enumerate the CFU after incubation in aerobic conditions at 30°C for 24 hours (Urbanek *et al*., 2020).

### 2.3. Selection of EPS-emulsifying degrading bacteria

Isolation of bacteria was performed from both the 10^−1^ diluted and undiluted intestinal content of each *T. molitor* larva. For the enrichment of the 10^−1^ dilution, a 0.5 mL aliquot was transferred to tubes containing 4.5 mL of carbon-free minimal basal broth and a 1 cm^2^ fragment of emulsified EPS. The tubes were incubated at 30°C for 7 days and at 25°C for 21 days (Tan *et al*., 2021b). Subsequently, aliquots from both the enriched and non-enriched dilutions were plated by dilution and streaking on nutrient agar and MacConkey agar. After incubation at 30°C for 48 hours, representative bacterial colony morphotypes were selected and subcultured on Tryptic Soy Agar (TSA) in duplicate, followed by Gram staining. Pure cultures were stored at 4°C.

To select EPS-degrading bacteria, they were cultured in duplicate in carbon-free minimal basal broth with a 1 cm^2^ fragment of emulsified EPS as the carbon source, and 1% glucose as a positive control. After incubation at 30°C for 7 days, the optical density of the cultures was measured using a visible light spectrophotometer (Model Tenso Med NV-203) at 600 nm, with non-inoculated EPS and glucose broth as the blank. The ten bacteria isolated on nutrient agar and the ten isolated on MacConkey agar that exhibited the highest optical density values in the minimal basal broth with EPS were selected. These were then plated in 5 mL of the same broth with EPS and 20 μL of a 1% solution of 2,3,5-triphenyltetrazolium chloride (TTC) as a viability indicator. The cultures were incubated at 30°C, with readings taken every 12 hours to observe any color change in the indicator to red, which would indicate metabolic activity and thus the utilization of EPS as a carbon and energy source (Tan *et al*., 2021b).

### 2.4. Comparison of the efficiency in the degradation of emulsified EPS

The efficiency of EPS degradation by the selected bacteria was determined indirectly by measuring the turbidity generated in the culture medium, which is related to microbial growth, and directly by determining the weight loss of the polymer or substrate consumed (Maroof *et al*., 2021), the reduction rate (k), and the half-life of the residual EPS (t_1_/_2_) (Auta *et al*., 2017). To determine turbidity, 3% of a 12-hour bacterial inoculum was added to 50 mL of minimal basal broth along with three 1 cm^2^ fragments of emulsified EPS. The mixture was incubated at 30°C, and its optical density at 600 nm was measured every 12 hours for 72 hours, comparing it with an uninoculated control and a control with glucose as the carbon source (Tang *et al*., 2017).

The weight loss of the polymer was determined by culturing the selected bacteria in triplicate flasks containing 100 mL of minimal basal medium and 0.36 – 0.48 g of emulsified EPS (initial weight). After incubation at 30°C for 100 days, the polymer was extracted and mixed with sodium dodecyl sulfate, then incubated at 50°C with constant agitation (40 rpm) to remove adhered bacterial cells (Wang *et al*., 2020). The polymer was then washed with distilled water and dehydrated at 60°C until a constant weight (final weight) was achieved. The percentage of weight loss was calculated based on this (Jiang *et al*., 2021).

The reduction rate (k) was calculated using the weight loss (Auta *et al*., 2017) (Equation 1):

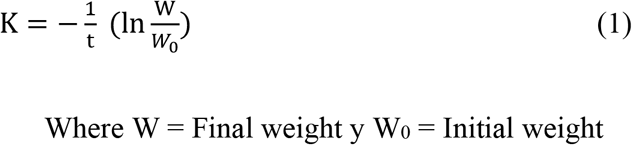

The half-life of the residual EPS (t_1_/_2_) (Auta *et al*., 2017) was calculated as follows (Equation 2):

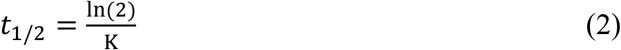

### 2.5. Molecular characterization of the microbial strain

Bacterial DNA extraction was carried out using the Wizard Genomic DNA Purification Kit (Promega). The 16S rRNA gene was amplified by PCR with the universal primers 27F and 1492R. The PCR products were then sequenced using the Sanger method at the “Marcel Gutiérrez Correa” Mycology and Biotechnology Laboratory at the Universidad Nacional Agraria La Molina (UNALM). The assembled sequences were preliminarily identified using BLASTN and aligned with related 16S rRNA gene sequences retrieved from the NCBI database (Altschul *et al*., 1997). Finally, phylogenetic analysis was performed using MEGA11 (Tamura *et al*., 2021), employing the Neighbor-Joining method and 1000 bootstrap replicates.

### 2.6. Data processing and statistical analysis

Bacterial efficiency in the degradation of PS, expressed as the reduction rate and half-life of residual PS, is presented as mean values with standard deviation (±). The mean weight loss due to the use of EPS as a carbon and energy source was analyzed using parametric statistics to determine differences between treatments (analysis of variance) and their significance (Tukey’s multiple comparison test), with a significance level of 0.05. IBM SPSS Statistics version 28 was used for the analysis.

## 3. RESULTS AND DISCUSSION

### 3.1. Number of cultivable bacteria associated with the intestines of *Tenebrio molitor* larvae

The range and percentage of cultivable bacteria associated with the intestines were from 1.0 × 10^5^ to 9.9 × 10^7^ CFU mL^-1^ in LCC (60%) and LRMD (65%), and from 1.0 × 10^5^ to 9.9 × 10^6^ CFU mL^-1^ in LREPS (85%) (Table 1, Figure 1).

**Table 1.**
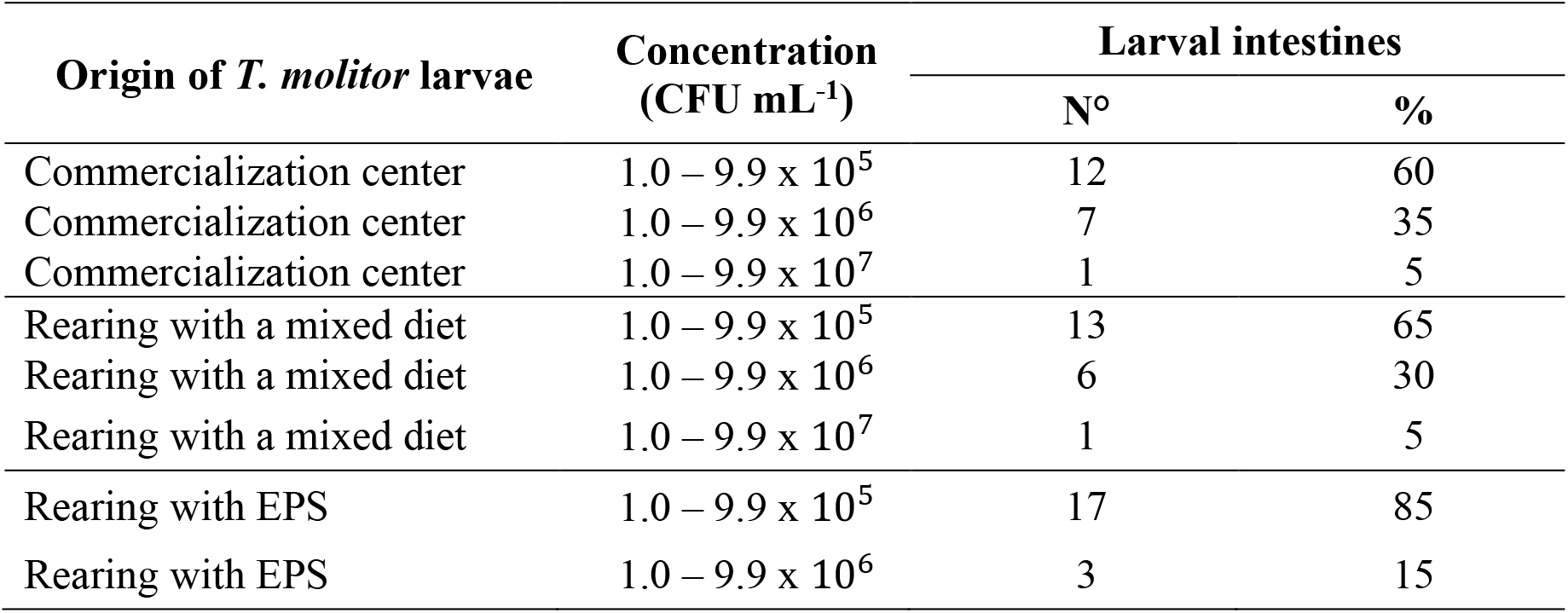
Concentration of cultivable bacteria (CFU/mL) associated with the intestines of *Tenebrio molitor* larvae.

| Origin of *T. molitor* larvae | Concentration (CFU mL<sup>-1</sup>) | Larval intestines |  |
| --- | --- | --- | --- |
|  |  | Nº | % |
| Commercialization center | $1.0 - 9.9 \times 10^5$ | 12 | 60 |
| Commercialization center | $1.0 - 9.9 \times 10^6$ | 7 | 35 |
| Commercialization center | $1.0 - 9.9 \times 10^7$ | 1 | 5 |
| Rearing with a mixed diet | $1.0 - 9.9 \times 10^5$ | 13 | 65 |
| Rearing with a mixed diet | $1.0 - 9.9 \times 10^6$ | 6 | 30 |
| Rearing with a mixed diet | $1.0 - 9.9 \times 10^7$ | 1 | 5 |
| Rearing with EPS | $1.0 - 9.9 \times 10^5$ | 17 | 85 |
| Rearing with EPS | $1.0 - 9.9 \times 10^6$ | 3 | 15 |

**Figure 1.**
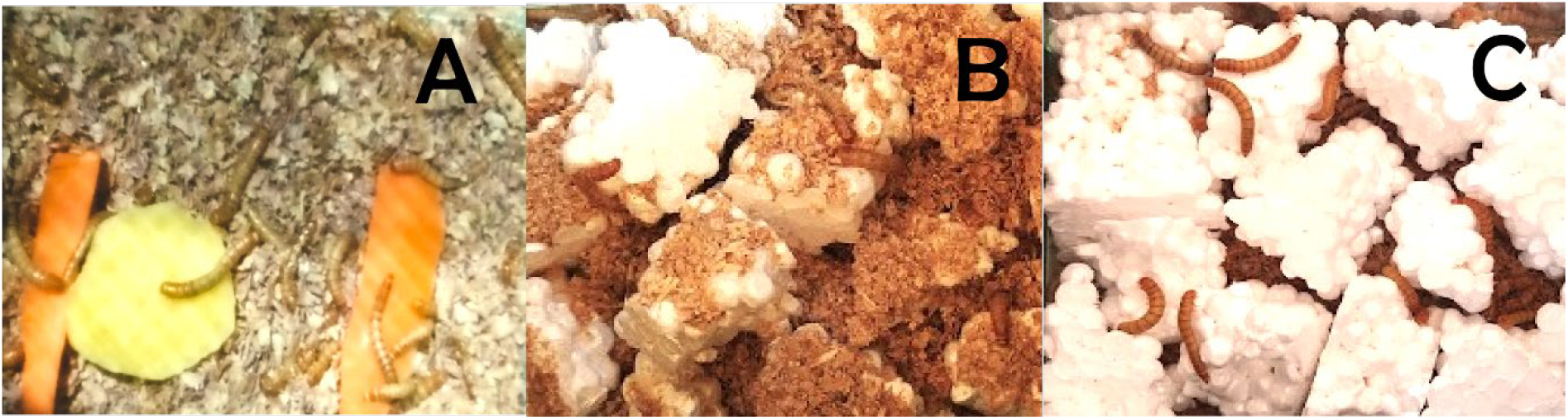
*T. molitor* larvae from the commercialization center (A), from laboratory rearing with a mixed diet of wheat bran and EPS (B), and from rearing with a diet consisting solely of EPS (C).

The hydrophobicity of most plastics, such as EPS, makes them resistant to hydrolysis, which negatively impacts their chemical or biological degradation (Ho *et al*., 2017). Despite this, *T. molitor* larvae from the Tenebrionidae family were able to consume EPS for 21 days, consistent with reports by Cunguan *et al*. (2023), Vera *et al*. (2021), and Brandon *et al*. (2021). EPS consumption was confirmed by the weight loss of the polymer, which indicates its use as a carbon source (Machona *et al*., 2022).

Microorganisms associated with the larvae’s intestines play a crucial role in the catabolism of EPS (Tsochatzis *et al*., 2021), without diminishing the larvae’s own activity in fragmenting the polymer into smaller particles (Machona *et al*., 2022). Brandon *et al*. (2021) reported no significant difference in the activity of the supernatant from the intestinal content of *T. molitor* larvae fed wheat bran or polystyrene (PS), both supplemented with antibiotics. In both cases, emulsifying activity was observed, suggesting that this property is intrinsic to the larvae’s intestines, regardless of the type of food consumed. Jiang *et al*. (2021) concluded that larvae and intestinal microorganisms have a symbiotic interaction. After ingesting plastic, intestinal microorganisms play a role in biodegradation, but the larvae’s enzymatic system also contributes to the degradation process.

The number of bacteria was lower in the intestines of *T. molitor* larvae fed EPS compared to larvae from the commercialization center and those fed a mixed diet. This result is consistent with the findings of Tsochatzis *et al*. (2021), who demonstrated that the number of bacteria in the intestines of *T. molitor* larvae fed a wheat bran: PS diet at a 4:1 ratio was lower than that of larvae fed a wheat bran diet (control). This difference was attributed to the bacteriostatic effect of PS degradation products, such as styrene and styrene oligomers. Additionally, Urbanek *et al*. (2020) showed that the highest number of colony-forming units of bacteria was found in the intestines of control larvae. This number varied depending on the type of PS, being higher with PS used for packaging (44.325 × 10^6^) compared to EPS, which has a molecular weight ratio (MFR) of polymer chain length 3.5 times greater.

### 3.2. Selected EPS-emulsifying bacteria

The range of colony morphotypes developed on nutrient agar was from one to four, resulting in 131 pure cultures from the non-enriched culture and 136 from the enriched culture. On MacConkey agar, the range of colony morphotypes was from one to five, yielding 102 pure cultures from the non-enriched culture and 114 from the enriched culture.

Among the 483 cultivable bacteria associated with the intestines of *T. molitor* larvae, 61.28% (296) were Gram-negative, while 38.72% (187) were Gram-positive. In basal minimal medium free of carbon with a fragment (1 cm^2^) of emulsified EPS, 95.13% (254) of the bacteria isolated on nutrient agar grew, and 86.57% (187) of the bacteria isolated on MacConkey agar grew. In total, 91.30% (441) of the bacteria grew with EPS, including 59.64% (263) Gram-negative and 40.36% (178) Gram-positive bacteria. From these, the 20 strains (ten from each agar type) with the highest optical density values were selected (Table 2).

**Table 2.** Selected bacteria with the highest optical density in basal minimal medium with emulsified EPS for degradation testing.

| Bacterial strains | Culture media | Optical density (600 nm) |  |
| --- | --- | --- | --- |
|  |  | Glucose (control) | Emulsified EPS* |
| 250 CIFOS | Agar Mc Conkey | 0.181 | 0.155 |
| 267 CIFOS | Agar nutritivo | 0.182 | 0.141 |
| 367 CIFOS | Agar nutritivo | 0.157 | 0.138 |
| 277 CIFOS | Agar nutritivo | 0.173 | 0.137 |
| 320 CIFOS | Agar nutritivo | 0.174 | 0.130 |
| 416 CIFOS | Agar nutritivo | 0.166 | 0.129 |
| 219 CIFOS | Agar Mc Conkey | 0.169 | 0.129 |
| 198 CIFOS | Agar Mc Conkey | 0.177 | 0.128 |
| 135 CIFOS | Agar nutritivo | 0.169 | 0.126 |
| 204 CIFOS | Agar Mc Conkey | 0.154 | 0.126 |
| 364 CIFOS | Agar nutritivo | 0.158 | 0.124 |
| 85 CIFOS | Agar Mc Conkey | 0.192 | 0.123 |
| 355 CIFOS | Agar nutritivo | 0.190 | 0.120 |
| 206 CIFOS | Agar Mc Conkey | 0.161 | 0.120 |
| 175 CIFOS | Agar nutritivo | 0.184 | 0.119 |
| 228 CIFOS | Agar Mc Conkey | 0.176 | 0.119 |
| 215 CIFOS | Agar Mc Conkey | 0.173 | 0.118 |

\*EPS = Expanded polystyrene

The isolation of bacteria from the intestines of *T. molitor* larvae is consistent with the findings of Machona *et al*. (2022). The enrichment technique was employed because it enhances the selection of microorganisms with greater efficiency in degrading PS (Brandon *et al*., 2021). The bacteria associated with the intestines of *T. molitor* larvae were predominantly Gram-negative, aligning with reports from Machona *et al*. (2022), Bae *et al*. (2021), and Peña *et al*. (2020). Additionally, it has been observed that the intestinal microbiome of *T. molitor* larvae is predominantly composed of the phyla Proteobacteria and Firmicutes, with very low percentages of Acidobacteria and Bacteroidetes (Urbanek *et al*., 2020).

Bacteria that grew using EPS as a carbon source (91.30%) were classified as polymer degraders. The biodegradation of plastics begins with microorganisms growing on the polymer surface, where they secrete enzymes to break it down into small fragments called oligomers and, eventually, into monomeric units. The pathways for the catabolism of styrene or EPS components are diverse; however, the most common involves the oxidation of styrene to phenylacetate via the citric acid cycle. Degradation products of EPS include styrene oxide, phenylacetaldehyde, phenylacetic acid, and phenylacetyl coenzyme A. The enzymes involved in this process are styrene monooxygenase (SMO), styrene oxide isomerase (SOI), phenylacetaldehyde dehydrogenase (PAALDH), and phenylacetyl coenzyme A ligase (PACoA) (Ho *et al*., 2017).

It is believed that PS-degrading bacteria can secrete extracellular oxidative enzymes that fragment PS chains and generate intermediates with C=O bonds. Identifying the key enzymes involved in the depolymerization and degradation of PS is crucial (Mamtimin *et al*., 2023); however, evidence suggests that EPS-degrading bacteria include Gram-negative members of the Enterobacteriaceae family and Gram-positive genera such as Enterococcus and Bacillus, among others (Sun *et al*., 2022; Tsochatzis *et al*., 2021; Tan *et al*., 2021a, 2021b). In environments where PS is the only carbon source, degrading bacteria can utilize the polymer for growth. However, if other carbon sources are present in addition to PS, the rate of contaminant degradation decreases. This limitation underscores the need for further research into the enzymes involved in the process and their potential for commercial application (Ho *et al*., 2017).

A reduction in the TTC indicator was observed in 65% (13) of the broths cultured with the bacteria (7 from nutrient agar and 6 from McConkey agar), as evidenced by the appearance of a stable raspberry color after 36–228 hours of incubation (Figure 2A). In the glucose control, the indicator reduction was observed between 24 and 72 hours. Among the TTC-reducing bacteria, the five strains that required 36 hours for TTC reduction while utilizing emulsified EPS as a carbon and energy source were selected.

**Figure 2.**
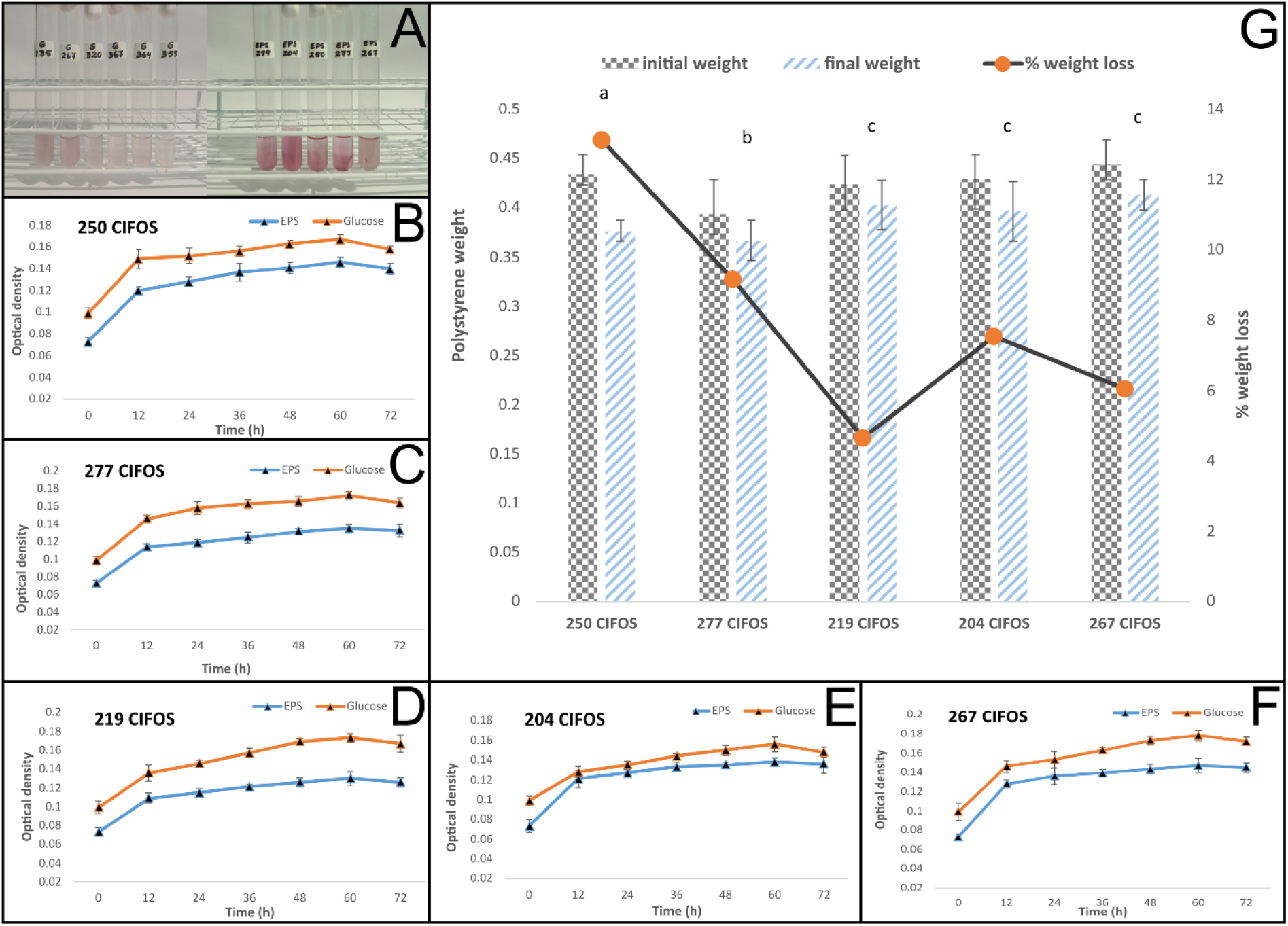
TTC Indicator (A), Optical density of minimal basal broth with glucose and EPS (B, C, D, E, F), and percentage of weight loss of EPS (G) utilized by intestinal bacteria from *Tenebrio molitor*.

The reduction of TTC by EPS-degrading bacteria was also demonstrated by Tan *et al*. (2021a) with bacteria isolated from the intestines of *Zophobas morio* larvae. The researchers observed TTC reduction from 24 to 96 hours of incubation, considered a primary screening for viability and metabolic activity. This rapid test verifies EPS degradation when used as a carbon source by the bacteria. The initially colorless TTC is reduced to the colored compound triphenylformazan (TPF) in the electron transport system. TTC is an artificial electron acceptor and is reduced in the aerobic cytochrome system to form the insoluble and colored compound called formazan (Tan *et al*., 2021b). The time required for TTC reduction by the intestinal bacteria of *T. molitor* larvae in the EPS broth (36–228 hours) was longer than in the broths with glucose as the carbon source (24–72 hours). This time difference was also observed by Tan *et al*. (2021b) with *B. megaterium* degrading PS, concluding that the polymer requires more time to be broken down into short monomers that can enter the bacterial cell and be used as a carbon source during growth.

### 3.3. Comparison of Bacterial Efficiency in the Degradation of Emulsified EPS

The optical density (600 nm) of the broths cultured with bacteria degrading emulsified EPS as a carbon and energy source gradually increased, reaching its maximum value at 60 hours. The timing was similar for bacteria cultured with the glucose control. At 60 hours, the optical density values ranged from 0.130 to 0.147 with EPS and from 0.156 to 0.178 with glucose. The highest optical density values were observed for bacterium 267, which utilized both EPS and glucose as carbon and energy sources (Figure 2B-G). Similarly, Lin and Liu (2021) evaluated the growth of intestinal bacteria from *T. molitor* and *Z. morio* larvae, using broth turbidity as an indicator of growth and PS degradation over 48 hours, and quantified CFUs to assess microbial viability.

The efficiency of EPS degradation, expressed as the percentage of weight lost by the degrading bacteria, ranged from 12.68% with bacterium 250 CIFOS to 5.29% with bacterium 267 CIFOS. Reported values in the literature vary, ranging from 7.40% to 12.97% (Jiang *et al*., 2021; Yang *et al*., 2015). EPS degradation by microorganisms is assessed through microscopic observation of changes in its surface structure (such as cracks, fissures, holes, and biofilm formation) and alterations in its mechanical and chemical properties (Cucini *et al*., 2022; Brandon *et al*., 2021; Tan *et al*., 2021a). Weight loss of the polymer (gravimetry) provides a direct measure of biodegradation; however, issues such as improper cleaning of the sample or weight loss due to volatilization of intermediate components and soluble impurities may affect the results (Ho *et al*., 2017).

The weight loss of the polymer degraded by bacterium 250 CIFOS reached 12.68%, which is higher than the 7.46% reported for *Bacillus cereus* by Auta *et al*. (2017). However, the reduction rate in this study was 0.0013 g/day with a half-life of 533.15 days (Table 3), compared to 0.0019 g/day and a residual polymer half-life of 363.16 days reported by Auta *et al*. (2017). This difference may be explained by the fact that Auta *et al*. (2017) used UV-irradiated PS for 25 days, a process that increases carbonyl groups and reduces tensile strength, thereby accelerating the rate of reduction and microbial degradation (Crystal *et al*., 2024).

**Table 3.** Reduction rate and half-life of residual EPS used as a carbon source by intestinal bacteria of *Tenebrio molitor*.

| Bacterial strains | Reduction rate (K)<br>(g days<sup>-1</sup>) | Half-life of residual EPS (t<sub>1/2</sub>)<br>(days) |
| --- | --- | --- |
| 250 CIFOS | 0.0013±0.0001 | 533.15±0.010 |
| 277 CIFOS | 0.0009±0.0002 | 770.11±0.034 |
| 204 CIFOS | 0.0006±0.0000 | 1155.17±0.125 |
| 267 CIFOS | 0.0006±0.0001 | 1155.17±0.009 |
| 219 CIFOS | 0.0005±0.0001 | 1386.20±0.145 |

The data presented are means of three replicates ± standard error.

### 3.4. Phylogenetic analysis

The BLASTN analysis identified the bacterium 250 CIFOS as belonging to the genus *Klebsiella*, while the phylogenetic analysis confirmed it as *K. pneumoniae* (GenBank accession number: PQ243593.1) (Figure 3).

**Figure 3.**
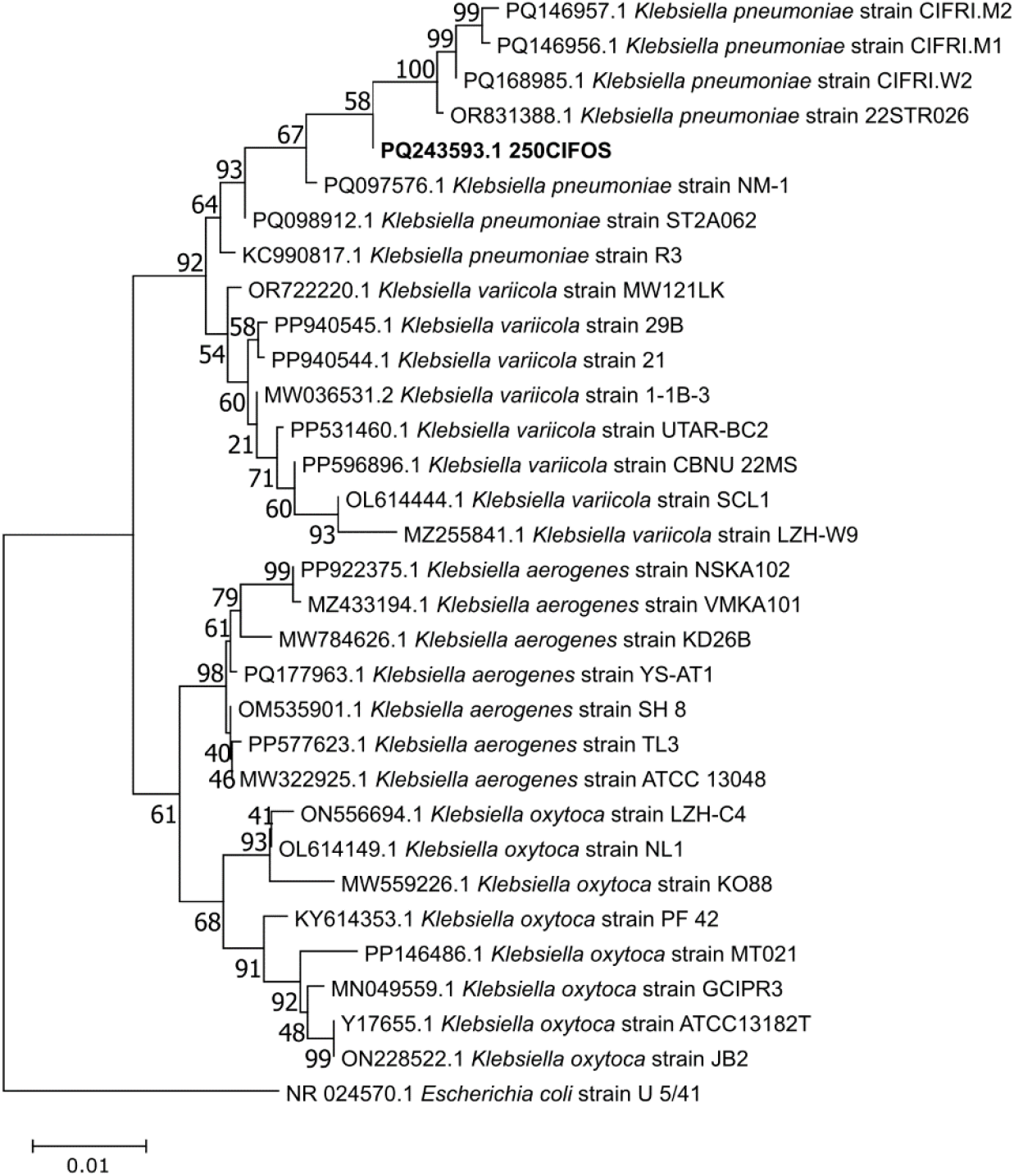
Phylogenetic tree of the 16S rRNA gene for strain 250 CIFOS and other *Klebsiella* sp. sequences, inferred using the Neighbor-Joining method with 1000 bootstrap replicates. The 16S rRNA sequence of *Escherichia coli* was used as the outgroup.

It has been demonstrated that bacteria of the genus *Klebsiella* (family Enterobacteriaceae) remained viable for 28 days in a broth with PS as the carbon source, with their adhesion and growth on the polymer confirmed by scanning electron microscopy (SEM) (Cucini *et al*., 2022). Additionally, the adhesion of *K. grimontii* to PS films, as verified by SEM, revealed a reduction in the size and sharpness of the film edges, as well as a deteriorated surface (Park *et al*., 2023). Furthermore, *K. oxytoca*, isolated from the intestines of *T. molitor* larvae that consumed PS, has been reported as a potential polymer degrader by Machona *et al*. (2022) and Urbanek *et al*. (2020), similar to *K. aerogenes* isolated from the intestines of *A. diaperinus* larvae (Cucini *et al*., 2022) and from the feces of *Z. morio* larvae (Sun *et al*., 2022), and *K. variicola* isolated from *Spodoptera frugiperda* larvae (Zhang *et al*., 2022). In this regard, Sun *et al*. (2022) concluded that emerging human pathogenic species such as *K. oxytoca, Enterococcus* spp., and *Corynebacterium* spp., identified in larvae that were starved and fed with PS, indicated that a diet with EPS is poor and induces an imbalance in the intestinal microbiome; however, these bacteria were associated with PS degradation. On the other hand, *K. pneumoniae* is an antibiotic-resistant enterobacteria (Araya *et al*., 2022), an opportunistic pathogen, and a causative agent of both nosocomial and community-acquired infections, especially in immunocompromised patients (Hartantyo *et al*., 2020). However, this soil-isolated bacterium has been reported as a degrader of microplastics (Saygin and Baysal, 2020) and polyethylene (Zhang *et al*., 2023). It was also recovered from the intestines of *T. molitor* larvae fed a diet of bran and PS (in a 4:1 ratio), along with the quantification (μg mg^-1^) of monomers and oligomers from PS degradation, such as acetophenone (0.180); 2,4-di-tert-butylphenol (0.100); 2,4,6-triphenyl-1-hexane (0.173); 1,1-diphenylethylene (0.016); and styrene (0.005) in the larvae feces (Tsochatzis *et al*., 2021).

## 4. CONCLUSIONS

Plastic-consuming larvae harbor several microorganisms with potential biotechnological applications. *Klebsiella pneumoniae* 250 CIFOS demonstrated the highest efficiency in degrading emulsified EPS, with a weight loss of 12.68%, a reduction rate of 0.0013 g/day, and a half-life of 533.15 days. While this bacterium is pathogenic, it serves as a valuable source of genes associated with enzymes that promote EPS degradation. Biotechnology offers the potential to accelerate and improve the efficiency of this biodegradation process.

## Notes

### Competing Interest Statement

The authors have declared no competing interest.

